## Supplemental figures and tables for "*Wolbachia* infection at least partially rescues the fertility and ovary defects of several new *Drosophila melanogaster bag of marbles* protein-coding mutants"

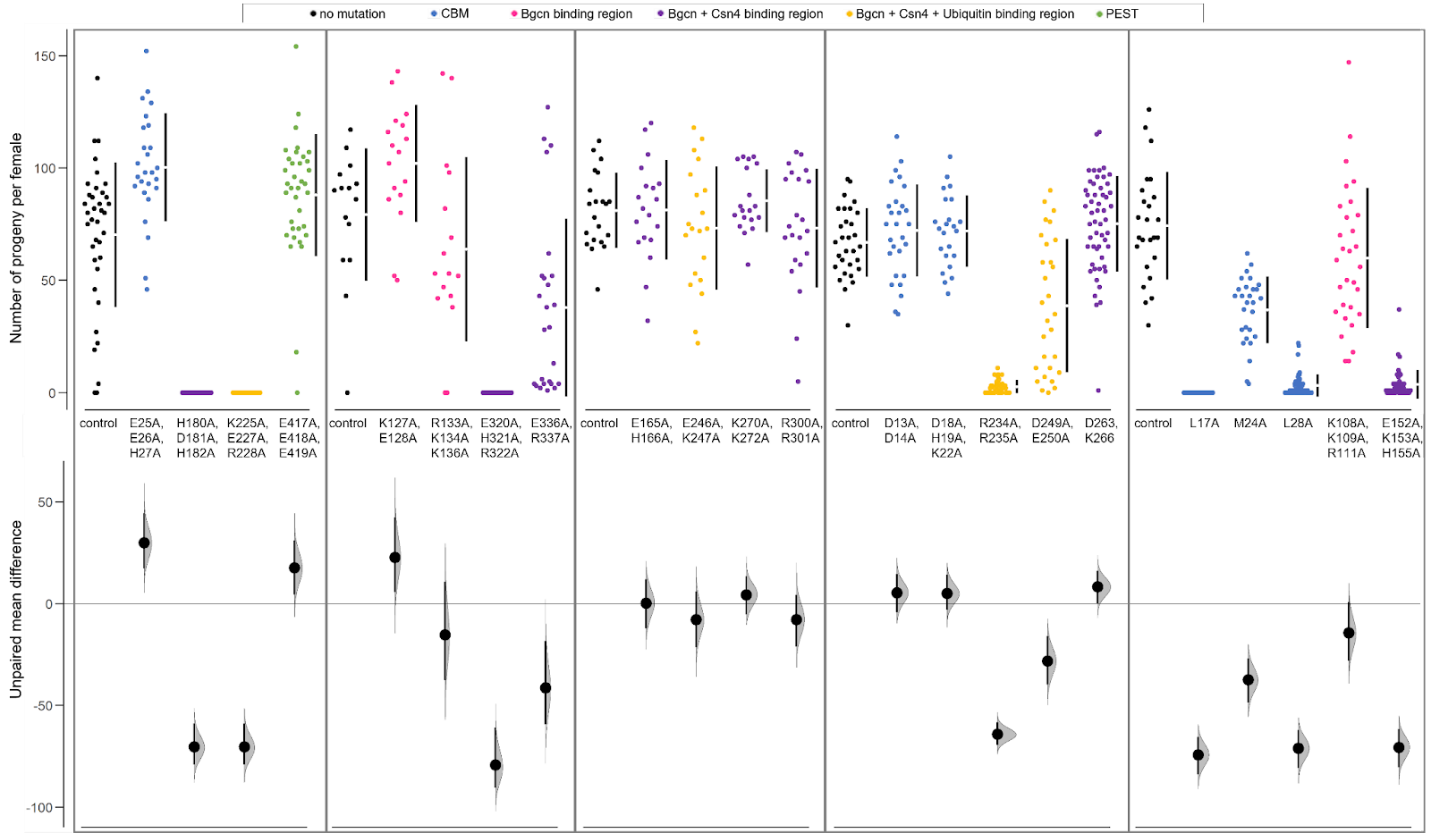

**Supplemental figure S1. Full estimation plots for fertility data of transgenic mutants compared to the *bam::Venus* transgenic control.** Fertility assays were done in five independent rounds. Below the jitter plots are the resampled bootstrap sampling distributions, with the mean differences represented by the black dots and the 95% confidence intervals represented by the vertical lines. Mean and sample size for each sample are listed in Supplemental Table S3.

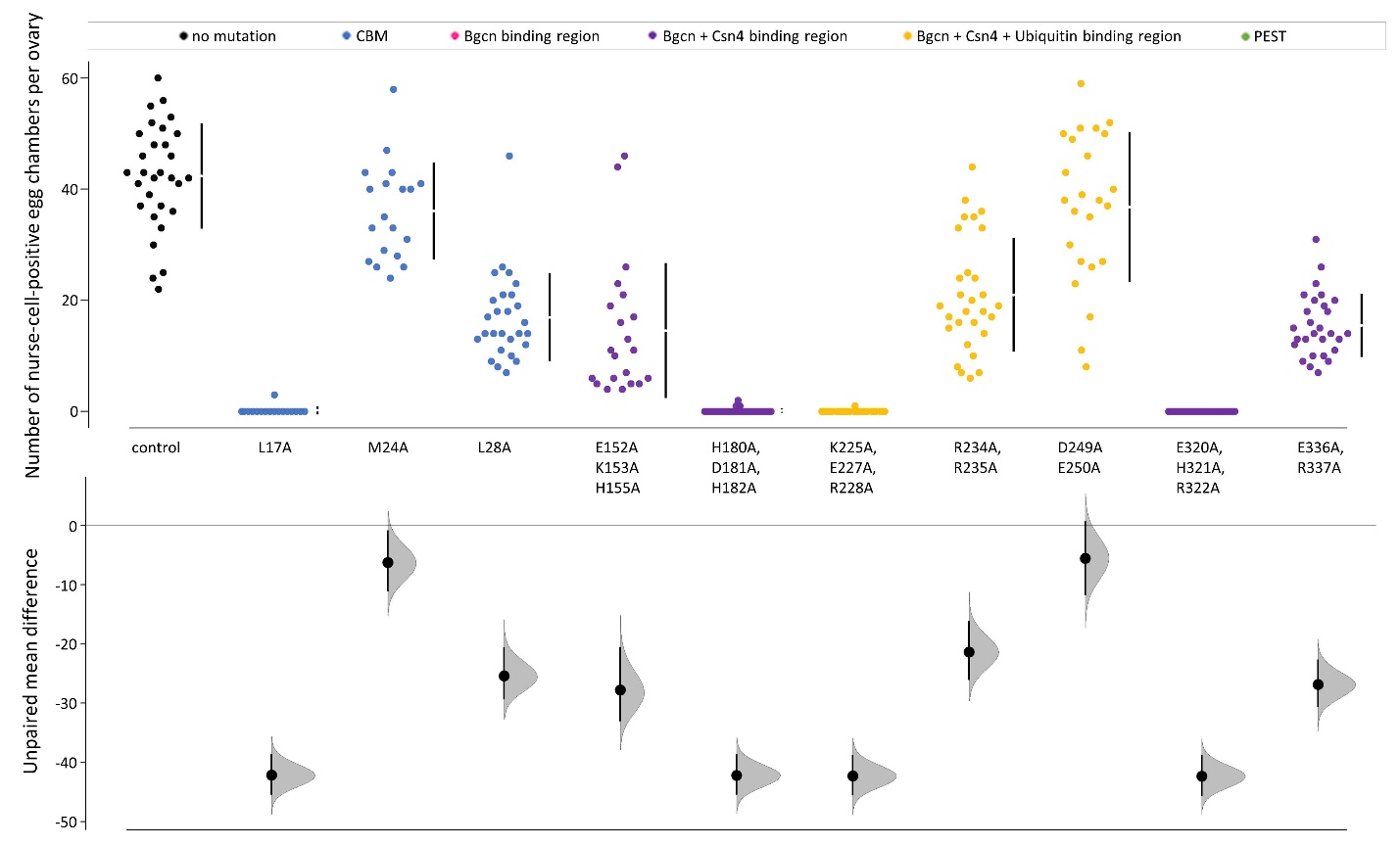

**Supplemental Figure S2. Full estimation plots for nurse-cell-positive egg chamber data of transgenic mutants compared to the *bam::Venus* transgenic control.** Below the jitter plots are the resampled bootstrap sampling distributions, with the mean differences represented by the black dots and the 95% confidence intervals represented by the vertical lines. Mean and sample size for each sample are listed in Supplemental Table S4.

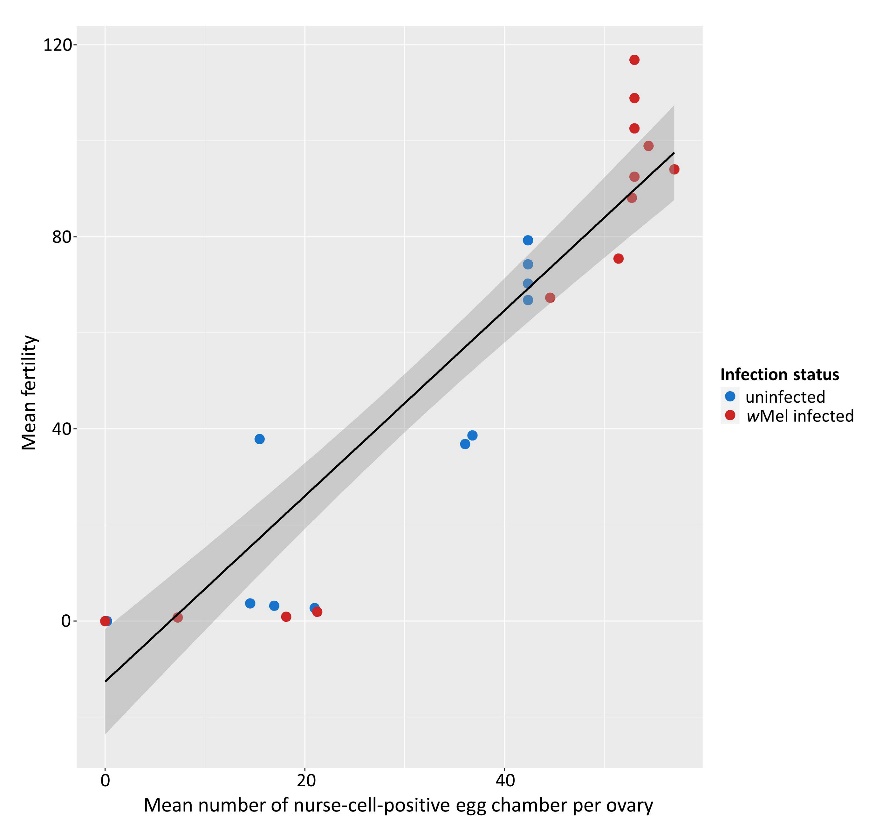

**Supplemental figure S3. Correlation of mean fertility and mean number of nurse-cell-positive egg chambers per ovary.** Each point represents the mean data for a fertility defective transgenic *bam* mutant line, with uninfected data in blue and infected data in red. Kendall’s rank correlation τ=0.828.

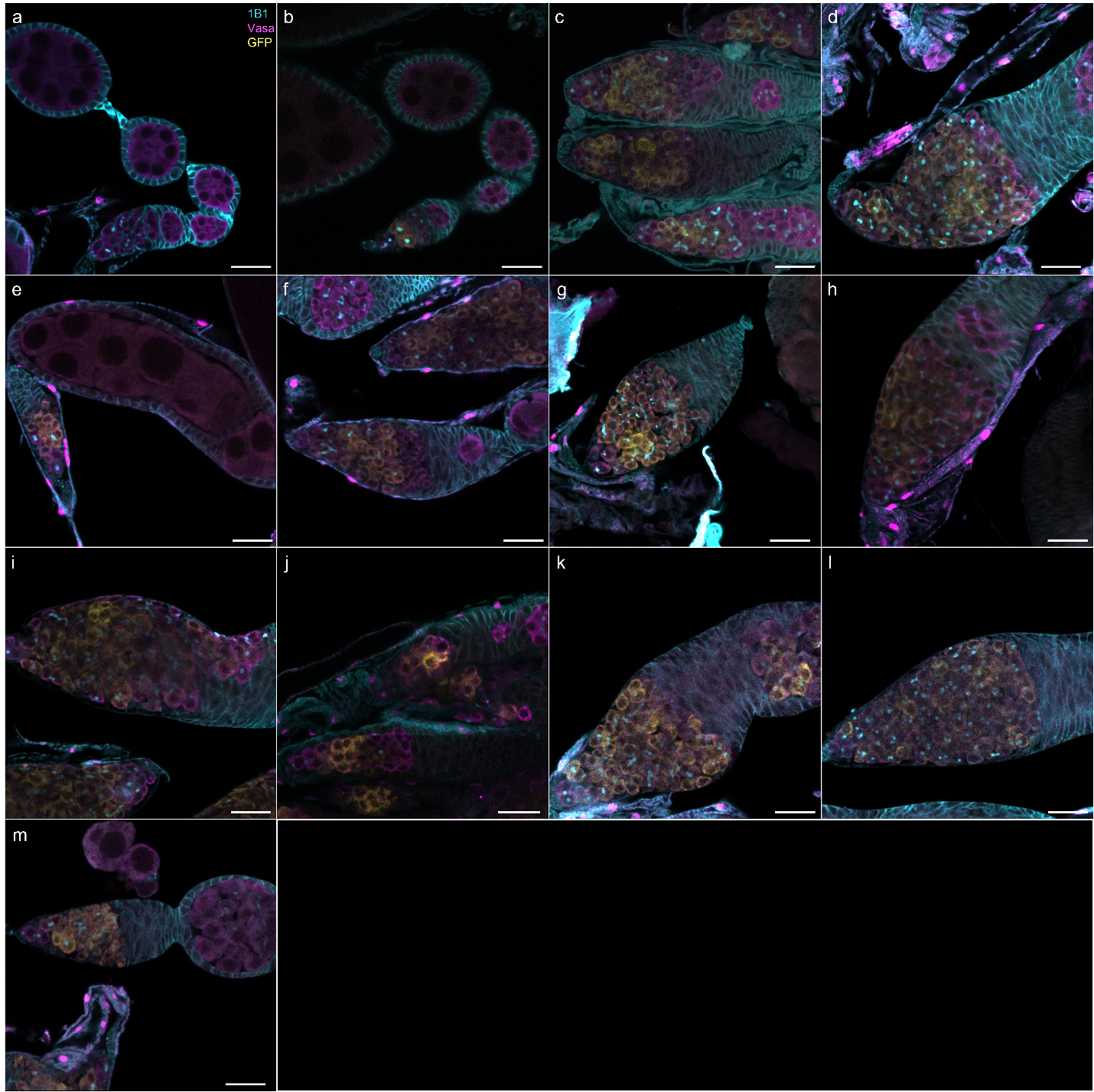

**Supplemental figure S4. Germariums of controls and all transgenic *bam* mutants with fertility defects when uninfected with *w*Mel.** Stained with anti-Hts-1B1 (cyan), anti-Vasa (magenta), and anti-GFP (yellow). (a) CantonS, (b) *bam::Venus* control, (c) *bam^L255F^::Venus*,
(d) *bam^L17A^::Venus*, (e) *bam^M24A^::Venus*, (f) *bam^L28A^::Venus*, (g) *bam^E152A, K153A, H155A^::Venus*,
(h) *bam^H180A, D181A, H182A:^::Venus*, (i) *bam^R234A, R235A^::Venus*, (j) *bam^D249A, E250A^::Venus*,
(k) *bam^K255A, E227A, R228A^::Venus*, (l) *bam^E320A, H321A, R322A^::Venus*, (m) *bam^E336A, R337A^::Venus*. Scale bar is 20μM for all images.

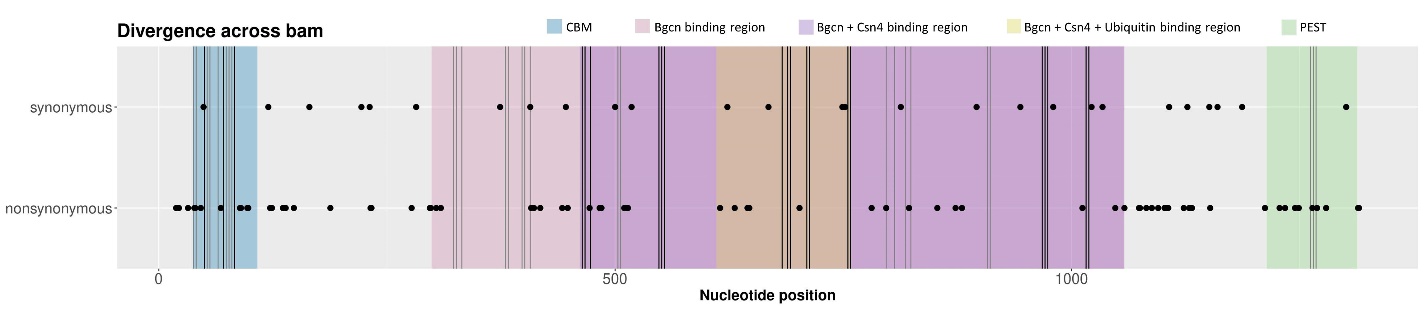

**Supplemental figure S5. Location of transgenic *bam* alanine mutants with respect to divergent amino acids between *D. melanogaster* and *D. simulans.*** Location of alanine mutations are represented by vertical lines, with fertility defect mutants in black and non-defect mutants in grey. Black dots represent the location of divergent amino acids between *D. melanogaster* and *D. simulans*, with synonymous divergences in the top row and nonsynonymous divergences in the bottom row. Background colors correspond to known functional and/or binding regions in *D. melanogaster*.

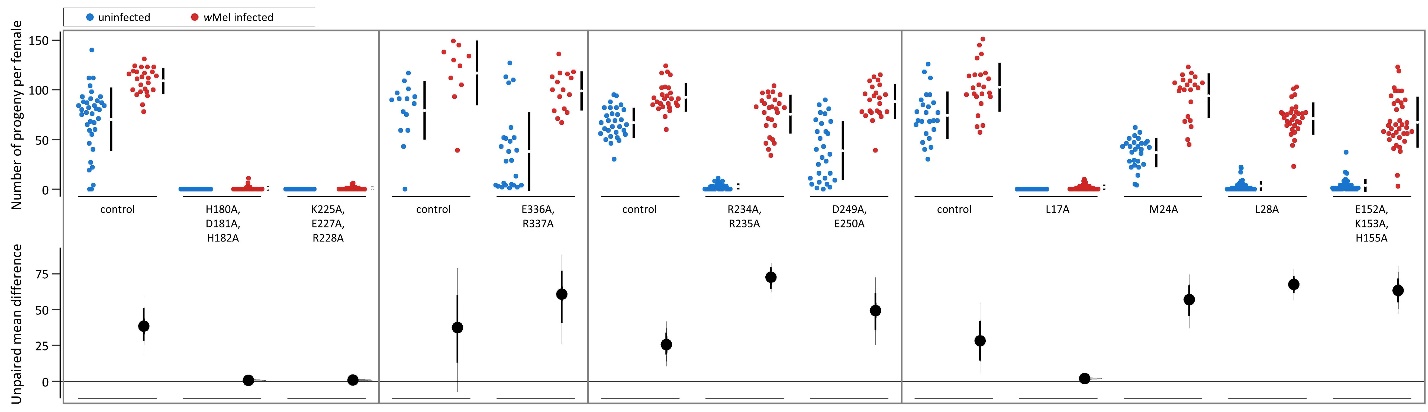

**Supplemental figure S6. Full estimation plots for fertility data of transgenic fertility-defective females, compared between uninfected and *w*Mel-infected.** Below the jitter plots are the resampled bootstrap sampling distributions, with the mean differences between the uninfected and infected data represented by the black dots and the 95% confidence intervals represented by the vertical lines. Mean and sample size for each sample are listed in Supplemental Table S3.

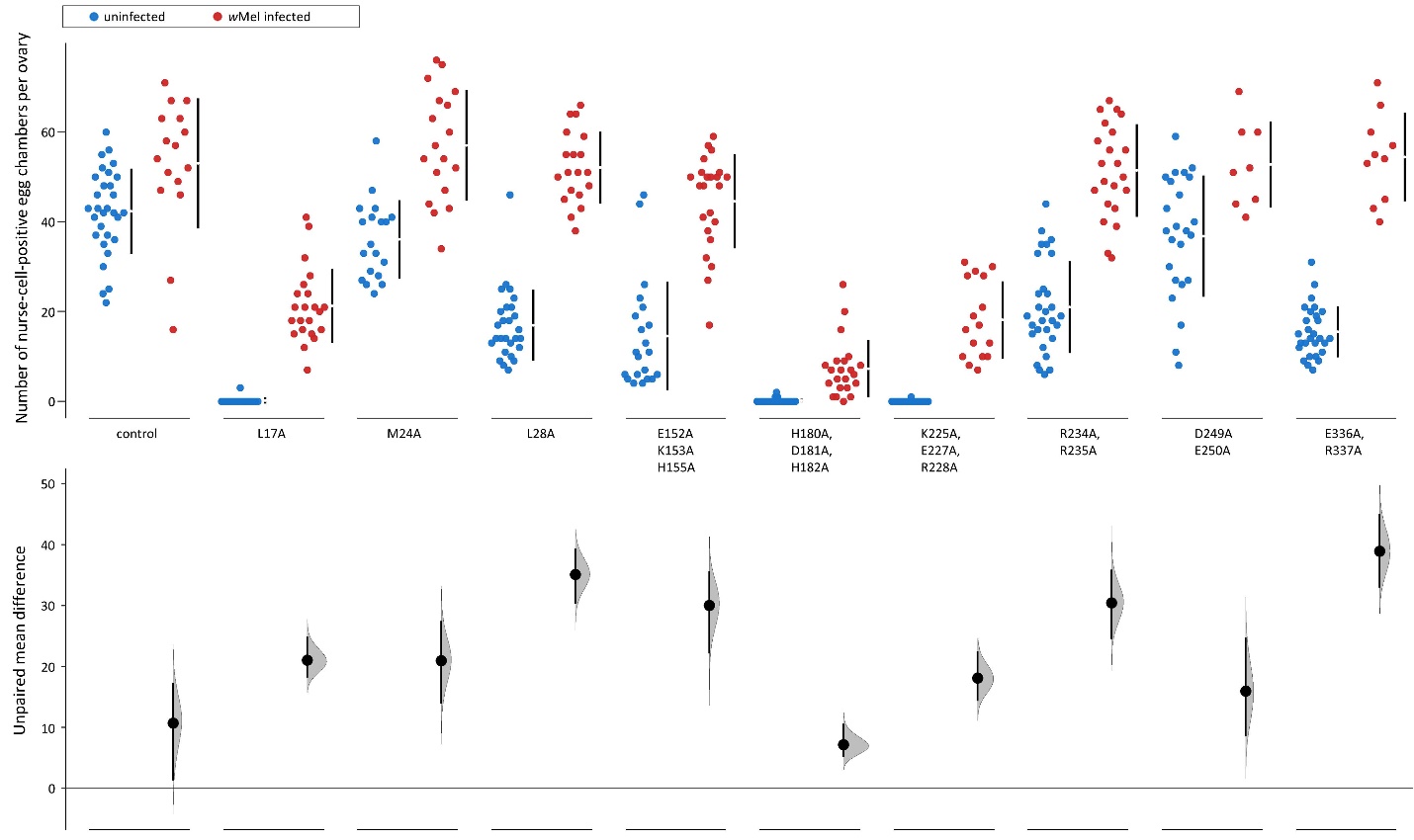

**Supplemental Figure S7. Full estimation plots for nurse-cell-positive egg chamber data of transgenic fertility-defective females, compared between uninfected and *w*Mel-infected.** Below the jitter plots are the resampled bootstrap sampling distributions, with the mean differences between the uninfected and infected data represented by the black dots and the 95% confidence intervals represented by the vertical lines. Mean and sample size for each sample are listed in Supplemental Table S4.

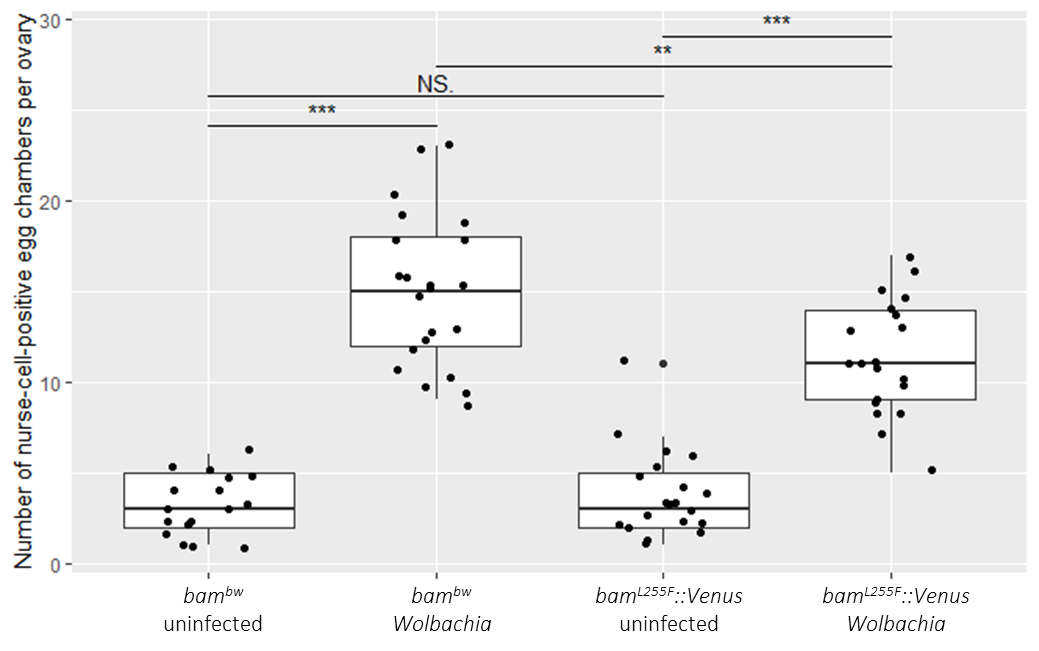
 **Supplemental figure S8. Comparison of original L255F mutant (*bam^bw^*) and the transgenic L255F mutant with a Venus tag (*bam^L255F^::Venus*) by the number of nurse-cell-positive egg chambers per ovary.** ** = significant at p<0.01; *** = significant at p<0.001; NS = not significant.

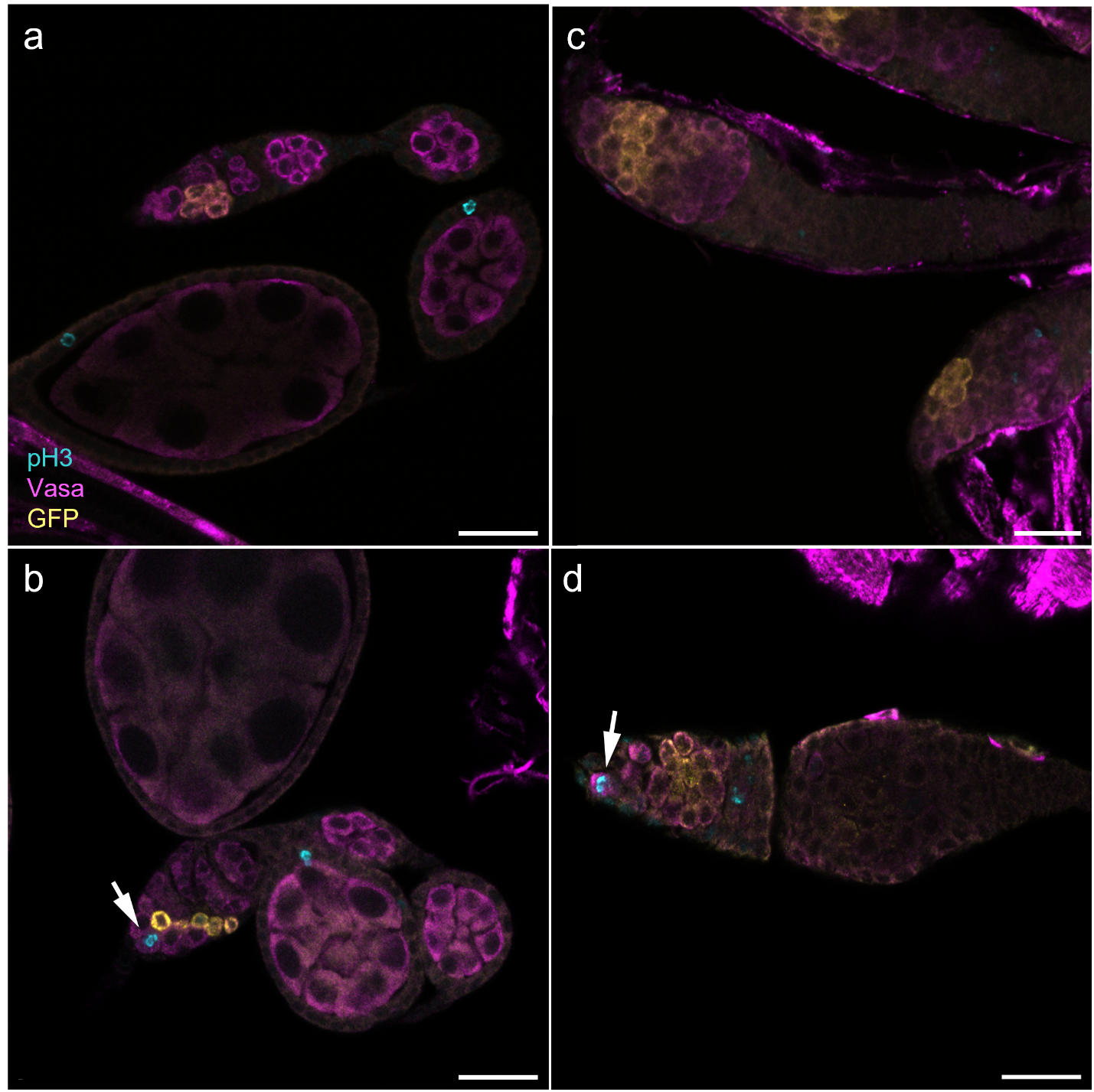

**Supplemental figure S9. GSC mitosis in the transgenic *bam::Venus* control and transgenic *bam^L255F^::Venus* hypomorph.** (a) No GSC mitosis and (b) active GSC mitosis in the *bam::Venus* control germarium. (c) No GSC mitosis and (d) active GSC mitosis in the *bam^L255F^::Venus* hypomorphic mutant.

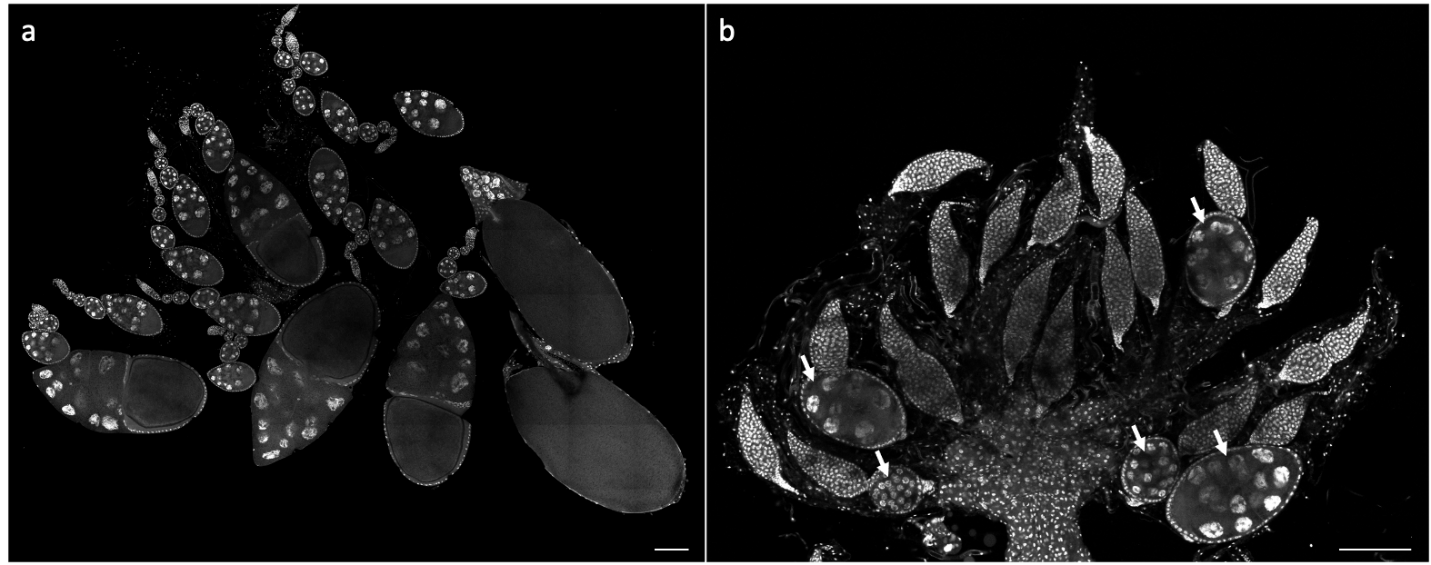

**Supplemental figure S10. Ovaries of a *bam* wildtype and *bam* RNAi knockdown fly.** (a) Ovaries of a *bam* wildtype fly. All egg chambers are results of successful Bam function. (b) Mutant ovaries of a nos-Gal4/+; UAS-*bam*^HMS00029^/+ uninfected fly. Arrows indicate nurse-cell-positive egg chambers that represent successful Bam function. Scale bar for both images is 100μM.

| **Species** | **Evidence of positive selection** | **Evidence for *Wolbachia*** |
| --- | --- | --- |
| *D. melanogaster* | Yes | Yes [54] |
| *D. simulans* | Yes | Yes [55] |
| *D. yakuba* | Yes | Yes [54, 56] |
| *D. santomea* | No | Yes [56] |
| *D. teissieri* | No | Yes [56] |
| *D. serrata* | Yes | No [57] |
| *D. bunnanda* | No | N/A |
| *D. birchii* | Yes | N/A |
| *D. jambulina* | Yes | No [57] |
| *D. bipectinata* | No | Yes [58]  No [59] |
| *D. pseudoananassae* | No | Yes [60]  No [59] |
| *D. pandora* | No | [61] |
| *D. ananassae* | Yes | Yes [54, 58] |
| *D. pseudoobscura* | No | Yes [58]  No [54, 57] |
| *D. affinis* | No | N/A |
| *D. mojavensis* | No | No [54, 57] |
| *D. immigrans* | No | No [57, 62] |
| *D. rubida* | Yes | No [60] |

**Supplemental table S1. Presence or absence of positive selection at *bam* and known *Wolbachia* infection status for *Drosophila* species.** Selection at *bam* is informed by McDonald-Kreitman tests from [30]. *Wolbachia* infection status for each species is cited individually in the table. N/A indicates there were no publications on infection status were found. Species that have evidence for both *Wolbachia* infection and absence reflect different stocks or geographical samples studied and were not included in the subsequent statistical analysis. In total, there are 4 species of *Wolbachia* infection and positive selection; 3 species of *Wolbachia* absence and positive selection; 3 species of *Wolbachia* infection and no positive selection; 2 species of *Wolbachia* absence and no positive selection. Fischer’s exact test results are p=1.0.

| **Bam mutations** | **Mutagenesis forward primer** | **Mutagenesis reverse primer** | **Annealing temp. (°C)** |
| --- | --- | --- | --- |
| D13A, D14A, | TGAGGGCAACgccgccCAGCAGTTGG | GGACACACGTCACGTGCATTAAG | 70 |
| L17A | CGACCAGCAGgctGACCACAATTTTAAGCAG | TCGTTGCCCTCAGGACAC | 65 |
| D18A, D19A, K22A, | ttttgccCAGATGGAGGAGCATTTGG | ttagcggcCAACTGCTGGTCGTCGTT | 64 |
| M24A | TTTTAAGCAGgctGAGGAGCATTTGGCC | TTGTGGTCCAACTGCTGG | 62 |
| E25A, E26A, H27A, | cgcTTTGGCCTTAATGGTGGAAG | gcggCCATCTGCTTAAAATTGTGG | 60 |
| L28A | GGAGGAGCATgctGCCTTAATGG | ATCTGCTTAAAATTGTGGTC | 58 |
| K108A, K109A, R111A, | tcggcaTCCGGAGGCGTGTTGGTC | ggcggcGGCCGGAGAACCGTCGAAT | 71 |
| K127A, E128A | GCAACTGCAGgccgctAATGTGTGGAACCGGAAGAGTAAAG | TTCTGCTTTGGCCCGGTG | 66 |
| R133A, K134A, K136A, | agtgctGGCTCTGCGTCCGCGGAT | ggctgcGTTCCACACATTTTCCTTCTGCAGTTGC | 71 |
| E152A, K153A, H155A | ctggcaATGATTGGTCTGCACGGC | agcggcAATAGTTATGGGCAGTTTCTCAATATTATC | 65 |
| E165A, H166A, | CAATAGCTTAgccgctAACGCCGTGC | TTTAGATAAGGAGCTTTTTATTATG | 58 |
| H180A, D181A, H182A, | cgctCTGACCGCCGATTTGGGC | gcagcCAGGGATCTGAACAGATTCATCAAAC | 66 |
| K225A, E227A, R228A | gccgctTTCCTTGTCCAGCAGCGC | tacagcGGTGCACAGCACCTGCAG | 69 |
| R234A, R235A | TGTCCAGCAGgctgctACCTTGGAGG | AGGAAGCGCTCTACTTTG | 61 |
| E246A, K247A, | CTTCGATTTCgccgctTACGACGAGTGTGACAAGTTG | TGGCGATTCGCCTCCAAG | 65 |
| D249A, E250A | gccgctTTCCTTGTCCAGCAGCGC | tacagcGGTGCACAGCACCTGCAG | 61 |
| L255F | TGACAAGTTGtTTAAGGGTTTC | CACTCGTCGTATTTCTCG | 58 |
| D263A, K266A, | ttcgctCTGCTTTTAAAGCCCAAAATG | gttggcCAAATAGGATGCGAAACC | 59 |
| K270A, K272A, | cgctATGCGCAATCGAAACGGA | ggggcTAAAAGCAGTTTGAAGTTGTCC | 62 |
| R300A, R301A, | AATTGGTCTGgctgccTGGATCAAGGCTGCGCATC | AGCAATCTCTCCATGCGC | 62 |
| E320A, H321A, R322A | tgctTACTCCGGGGCCATGACC | gcggcCAGATCCATTTCCCAGTTAAATACGTG | 67 |
| E336A, R337A | GTCGTTGAACgccgcaGCCATCCTTTTGTCC | TTGTGGCTTTCGGTCATG | 59 |
| E417A, E418A, E419A. | cgccTTTGAGGAGACCGAGGAAGTGC | gcggcCGACGCTGACGGCTGCTC | 70 |

**Supplemental table S2. Mutagenesis primers used to generate new *bam* mutants.** Each row lists the corresponding primer sequences and annealing temperature used in the mutagenesis PCR step of the cloning procedure to generate the respective amino acid mutant.

| **Transgenic *bam* allele** | **Uninfected mean fertility** | **Infected mean fertility** | **Uninfected sample size** | **Infected sample size** | **Assay round** |
| --- | --- | --- | --- | --- | --- |
| *bam^D13A, D14A^::Venus* | 72.2142857 | 90.3571429 | 28 | 28 | two |
| *bam^L17A^::Venus* | 0.0000000 | 1.9600000 | 43 | 50 | five |
| *bam^D18A, H19A, K22A^::Venus* | 71.9130435 | 90.8421053 | 23 | 19 | one |
| *bam^M24A^::Venus* | 36.8928571 | 94.0909091 | 28 | 22 | five |
| *bam^E25A, E26A, H27A^::Venus* | 100.2307692 | 102.0645161 | 26 | 31 | one |
| *bam^L28A^::Venus* | 3.2000000 | 71.0000000 | 50 | 31 | five |
| *bam^K108A, K109A, R111A^::Venus* | 59.9333333 | 98.6800000 | 30 | 25 | five |
| *bam^K127A, E128A^::Venus* | 102.0000000 | 119.1111111 | 17 | 9 | three |
| *bam^R133A, K134A^::Venus* | 63.8750000 | 107.5000000 | 16 | 10 | three |
| *bam^E152A, K153A, H155A^::Venus* | 3.6666667 | 67.3421053 | 45 | 38 | five |
| *bam^E165A, H166A^::Venus* | 81.3684211 | 100.8000000 | 19 | 10 | four |
| *bam^H180A, D181A, H182A^::Venus* | 0.0000000 | 0.7714286 | 34 | 35 | one |
| *bam^K225A, E227A, R228A^::Venus* | 0.0000000 | 0.9200000 | 29 | 25 | one |
| *bam^R234A, R235A^::Venus* | 2.7058824 | 75.4814815 | 34 | 27 | two |
| *bam^E246A, K247A^::Venus* | 73.2500000 | 104.8888889 | 20 | 18 | four |
| *bam^D249A, E250A^::Venus* | 38.6896552 | 88.173913 | 29 | 23 | two |
| *bam^D263A, K266A^::Venus* | 75.1320755 | 88.1111111 | 53 | 45 | two |
| *bam^K270A, K272A^::Venus* | 85.4210526 | 99.5384615 | 19 | 13 | four |
| *bam^R300A, D301A^::Venus* | 73.2727273 | 102.4583333 | 22 | 24 | four |
| *bam^E320A, H321A, R322A^::Venus* | 0.0000000 | 0.0000000 | 41 | 39 | three |
| *bam^E336A, R337A^::Venus* | 37.8800000 | 98.9375000 | 25 | 16 | three |
| *bam^E417A, E418A, E419A^::Venus* | 87.9166667 | 111.4347826 | 36 | 23 | one |
| *bam::Venus* | 70.2702703 | 108.9200000 | 37 | 25 | one |
| *bam::Venus* | 66.8666667 | 92.5384615 | 30 | 26 | two |
| *bam::Venus* | 79.2666667 | 116.9000000 | 15 | 10 | three |
| *bam::Venus* | 81.0952381 | 101.1875000 | 21 | 16 | four |
| *bam::Venus* | 74.2800000 | 102.6086957 | 25 | 23 | five |

**Supplemental table S3. Mean adult progeny per female seven days after first progeny eclosed, as uninfected and infected with *w*Mel.** *Bam* alleles are listed in order of location of mutations from N- to C-terminal, with mean fertility of and sample size of surviving females at the end of the assay. The corresponding assay in which the transgenic flies were evaluated is indicated by the fertility assay round.

| **Transgenic *bam* allele** | **Uninfected mean nurse-cell positive egg chambers per ovary** | **Infected mean nurse-cell positive egg chambers per ovary** | **Uninfected sample size** | **Infected sample size** |
| --- | --- | --- | --- | --- |
| *bam^L17A^::Venus* | 0.1764705882 | 21.22727273 | 17 | 22 |
| *bam^M24A^::Venus* | 36.05263158 | 57.0 | 19 | 18 |
| *bam^L28A^::Venus* | 16.92592593 | 52.05263158 | 27 | 19 |
| *bam^E152A, K153A, H155A^::Venus* | 14.52380952 | 44.56521739 | 21 | 23 |
| *bam^H180A, D181A, H182A^::Venus* | 0.1315789474 | 7.272727273 | 38 | 22 |
| *bam^K225A, E227A, R228A^::Venus* | 0.03703703704 | 18.125 | 27 | 16 |
| *bam^R234A, R235A^::Venus* | 20.96551724 | 51.40909091 | 29 | 21 |
| *bam^D249A, E250A^::Venus* | 36.79166667 | 52.75 | 24 | 8 |
| *bam^E320A, H321A, R322A^::Venus* | 0 | 0 | 33 | 24 |
| *bam^E336A, R337A^::Venus* | 15.46428571 | 54.4 | 28 | 10 |
| *bam::Venus* | 42.33333333 | 53 | 30 | 16 |

**Supplemental table S4. Mean number of nurse-cell-positive egg chambers per ovary, as uninfected and infected with *w*Mel.** *Bam* alleles are listed in order of location of mutations from N- to C-terminal, with mean number of nurse-cell-positive egg chambers per ovary and sample size.
